# Modelling Inoculum Availability of *Plurivorosphaerella nawae* in Persimmon Leaf Litter with Bayesian Beta Regression

**DOI:** 10.1101/771667

**Authors:** Joaquín Martínez-Minaya, David Conesa, Antonio López-Quílez, José Luis Mira, Antonio Vicent

## Abstract

Circular leaf spot (CLS), caused by *Plurivorosphaerella nawae*, is a serious disease of persimmon (*Diospyros kaki*) inducing necrotic lesions on leaves, defoliation and fruit drop. Under Mediter-ranean conditions, *P. nawae* forms pseudothecia in the leaf litter during winter and ascospores are released in spring infecting susceptible leaves. Persimmon growers are advised to apply fungicides for CLS control during the period of inoculum availability, which was defined based on ascospore counts under the microscope. A model of inoculum availability of *P. nawae* was developed and evaluated as an alternative to ascospore counts. Leaf litter samples were collected weekly in L’Alcúdia from 2010 to 2015. Leaves were soaked, placed in a wind tunnel, and released ascospores of *P. nawae* were counted. Hierarchical Bayesian beta regression methods were used to fit the dynamics of ascospore production in the leaf litter. The selected model, having the lowest values of DIC, WAIC and LCPO, included accumulated degree days (*ADD*) and *ADD* taking into account the vapor pressure deficit (*ADDvpd*) as fixed effects, and year as a random effect. This model had a MAE of 0.042 and RMSE of 0.062. The beta regression model was evaluated in four orchards for different years from 2010 to 2015. Higher accuracy was obtained at the beginning and the end of the ascospore production period, which are the events of interest to schedule fungicide sprays for CLS control in Spain. This same modelling framework can be extended to other fungal plant pathogens whose inoculum dynamics are expressed as proportion data.

## 1 Introduction

Circular leaf spot (CLS) disease of persimmon (*Diospyros kaki* Thunb.), caused by *Plurivo-rosphaerella nawae* (Hiura & Ikata) O. Hassan & T. Chang (= *Mycosphaerella nawae*), induces necrotic lesions on leaves, chlorosis and defoliation. The presence of foliar lesions and premature leaf drop induce early fruit maturation and abscission, resulting in serious economic losses (Bassimba et al., 2017). The disease was first described in humid areas in Japan and Korea (Ikata and Hitomi, 1929; Kang et al., 1993). The detection of CLS in Eastern Spain was the first report of the disease in a semi-arid area (Vicent et al., 2012).

The fungus forms pseudothecia in leaf litter during winter and ascospores are produced as temperatures increase in spring (Kang et al., 1993). Ascospores are wind-dispersed and infect persimmon leaves in the presence of a film of water and adequate temperatures. The main infection period in Korea was from mid-May to the end of July (Kang et al., 1993; Kwon and Park, 2004) and from the beginning of April to early July in Spain (Vicent et al., 2012). The asexual stage of *P. nawae* was identified in Korea as belonging to the genus *Ramularia*, but its role in field epidemics is not fully understood (Kwon et al., 1998; Kwon and Park, 2004). In Spain, this secondary inoculum has not been observed (Vicent et al., 2012). The disease is characterised by a long incubation period of about 4 months (Kwon and Park, 2004; Vicent et al., 2012).

Fungicide schedules for the control of CLS in Korea consist of three to four foliar applications during the infection period. Although the efficacy of fungicide programs may differ depending on the year, good disease control was obtained under experimental conditions (Kwon et al., 1998; Kwon and Park, 2004). In Spain, two to four fungicide applications during the infection period in spring showed also good efficacy for the control of CLS, whereas post-infection sprays were ineffective (Bassimba et al., 2017; Berbegal et al., 2013). Cultural practices, such as leaf litter removal and moving from flood to drip irrigation systems, are also recommended to growers, but their efficacy has not been quantified so far (Vicent et al., 2011, 2012).

Fungicide programs are effective for CLS control only when spray applications coincide with the infection period, defined by the presence of ascospores, adequate environmental conditions and susceptible leaves. The presence of airborne ascospores is typically monitored using spore traps, either active volumetric or passive (West and Kimber, 2015). Nevertheless, the predictive ability of spore traps is somehow limited because they only detect the ascospores when already released in the orchard air. In the case of *P. nawae* in Spain, monitoring inoculum production in the leaf litter allowed to predict ascospore release 1–2 weeks in advance, so this method is routinely used by the advisory services to schedule fungicide sprays for CLS control (Vicent et al., 2012). Samples of leaf litter are collected weekly in affected persimmon orchards and soaked in distilled water. Immediately after soaking, leaves are placed in a wind tunnel until they dry. Ascospores released from the leaf litter are collected on glass microscope slides and counted under the microscope (Vicent et al., 2011). Although this method proved to be useful, it is time and resource consuming, requires specific laboratory equipment and qualified personnel. Consequently, the extent of the monitored area and the density of the sampling network are rather limited.

Models for inoculum maturation in the leaf litter have been developed for several ascomycetes, as a more efficient alternative to ascospore counts (De Wolf and Isard, 2007). Most of these models rely on transformations of the response variable and then fitted as a linear regression (Luley and McNabb Jr, 1991; Spotts et al., 1994). For instance, Gadoury et al. (1982) used a linear regression with a probit transformation to the proportion of ascospore discharge. Villalta et al. (2001) depicted a linear regression with a logit transformation. Rossi et al. (2009) and Eikemo et al. (2011) compared linear regressions with asymptotic, monomolecular, logistic and Gompertz transformations. In other cases, nonlinear regression was used (Navas-Cortés et al., 1998b; Rossi et al., 1999; Cooley et al., 2007; Legler et al., 2014). Nevertheless, as the proportion of ascospores is the variable being modeled, there are other methods available such as the beta regression model, firstly introduced by Ferrari and Cribari-Neto (2004). Basically, this methodology consists on assuming that the response variable conditioned to the linear predictor follows a beta distribution which is depending on two parameters, a mean and a precision.

On the other hand, Bayesian hierarchical methods are becoming popular in many fields as they better address the intrinsic complexity typical in many natural systems (Clark, 2005). In Bayesian inference, parameters are treated as random variables and data are related to model parameters using a likelihood function, getting the posterior distribution by combining the prior distribution and the likelihood function. However, getting the posterior distribution is not always straightforward and numerical algorithms are usually required. Markov Chain Monte Carlo (MCMC) methods (Gilks et al., 1996) are widely used to obtain posterior distributions but they involve computationally and time intensive simulations. The Integrated Nested Laplace Approximation (INLA) approach was developed as a computationally efficient alternative to MCMC in latent Gaussian models (Tierney and Kadane, 1986; Rue et al., 2009).

In this work, we used hierarchical Bayesian beta regression models with fixed and random effects to estimate the production *P. nawae* ascospores in persimmon leaf litter with the INLA methodology. The resulting model will assist to predict the dynamics of *P. nawae* inoculum in the leaf litter based on environmental covariates, without the direct quantification of ascospores in the leaf litter. This will facilitate a wider implementation of a decision support system to optimize the fungicide programs for CLS control in Spain.

## 2 Materials and Methods

### 2.1 Field data

The model was developed from 2010 to 2015 in a persimmon cv. Rojo Brillante orchard of 0.83 ha severely affected by CLS at L’Alcúdia in Valencia Province, Spain. Trees were 11 yr old at the beginning of the study and were grafted on *D. lotus* L. rootstock. The orchard was drip irrigated and had a north-south row orientation with a 5 m across-row spacing and 4 m in-row spacing. Orchards of similar characteristics were selected in Valencia Province at Benimodo, Villanueva de Castellón and Guadassuar (2010 and 2011) and Moncada (2012 to 2015) for model evaluation (i.e. validation). In all cases, an experimental area of 0.2 hectares (10 × 10 trees) in the center of each orchard remained without fungicide applications during the period of study.

Environmental data were monitored hourly in each orchard with an automated meteorological station (Hobo U30, Onset Computer Corp.) including sensors for temperature and relative humidity (Hobo S-THB, accuracies ± 0.2°C, ± 2.5%), and rainfall (7852, Davis Instruments Corp, resolution 0.2 mm). Environmental monitors were located at 1.5 m above the soil surface within the row in the center of the experimental area.

Following Rossi et al. (2009) time was expressed in physiological units calculated by three different methods, all of them based on sums of the daily mean air temperatures exceeding 0°C. In particular, accumulated degree days (*ADD*) were calculated as:

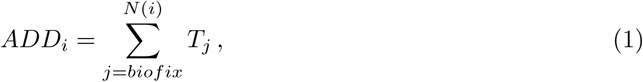

where *i* and *j* are the subscripts for observations and days, respectively, with *j* = *biofix* to *N* (*i*), while *T*_*j*_ is the air temperature in each day (calculated as the mean of 24 hourly values) if *T*_j_ > 0, elsewhere *T*_*j*_ = 0. The biofix was set at 1 January.

In second place, *ADD* considering vapor pressure deficit (*ADDvpd*) were calculated:

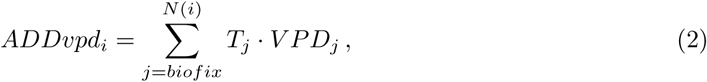

being *i* the observation and *j* the subscript for days, with *j* = *biofix* to *N* (*i*), and *T*_*j*_ is the air temperature in each day (calculated as the mean of 24 hourly values) if *T*_j_ > 0, elsewhere *T*_*j*_ = 0. *V PD_j_* is a dichotomic variable calculated as follows: when vapor pressure deficit (*vpd*)_*j*_ ≤ 4*hPa, V PD_j_* = 1, elsewhere *V* PD_*j*_ = 0, being *vpd* calculated from temperature an relative humidity (*rh*, %) as follows:

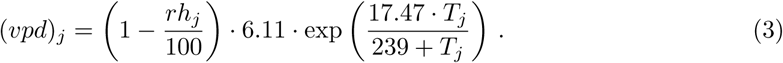

Finally, a variable that incorporates information about rainfall was also considered. In particular, it was denoted as *ADDwet* and corresponds to the accumulated degree days but taking into account both the *vpd* and rainfall (*R*):

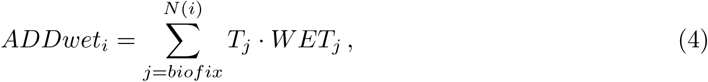

being *i* the observation and *j* the subscript with *j* = *biofix* to *N* (*i*), and *T*_*j*_ is the air temperature in each day (calculated as the mean of 24 hourly values) if *T*_j_ > 0, elsewhere *T*_*j*_ = 0. *WET*_*j*_ is a dichotomic variable calculated as follows: when *R*_*j*_ *≥* 0.2mm and *vpd*_*j*_ *≤* 4*hPa, WET_j_* = 1, elsewhere *WET*_*j*_ = 0.

The dynamics of *P. nawae* ascospore production in the leaf litter was studied in the orchards and years indicated above. Dry leaves on the orchard floor were covered with a plastic mesh (2 × 2 m^2^, 5-by-5-mm openings) fixed with four stainless-steel pins. Plastic nets were located in the center of the experimental area in each orchard without overlying the soil area wetted by the drip irrigation system. Leaf litter density under the plastic nets was adjusted to 350 g of dry leaves m^2^ (Vicent et al., 2011). A pooled sample of 20 dry leaves was collected weekly in each orchard, but four samples of 20 dry leaves were collected at L’Alcúdia from 2013 to 2015. Leaf litter samples were soaked for 15 min in distilled water. Immediately after soaking, leaves were placed with the abaxial surface facing upward in a wind tunnel for 30 min until they were visibly dry (Whiteside, 1974; Vicent et al., 2011). During the process, air and water temperature was maintained at about 21°C.

Discharged ascospores were collected on a glass microscope slide (26 × 76 mm) coated with silicone oil (Merck). Spores were stained with lactophenol-acid cotton blue and examined at 400X magnification. All ascospores showing the morphological characteristics of *P. nawae*; spindle-shaped, 10 *−* 13 × 3 *−* 4 *µ*m, hyaline, 2-celled with a medium or slightly supramedian septum (Kwon et al., 1998), were counted in four microscope field transects. Isolations were arbitrarily performed each year using additional leaf litter samples and collecting the ejected ascospores in potato dextrose agar (PDA) amended with 0.5 g L^−1^ of streptomycin sulphate (PDAS). Identification of the resulting fungal colonies was confirmed using a specific molecular method for *P. nawae* (Berbegal et al., 2013). For each week, the cumulative proportion of ascospores discharged was calculated based on the total collected in each orchard and year.

### 2.2 Beta regression

Beta regression is commonly used for variables that assume values in the unit interval (0,1) (Ferrari and Cribari-Neto, 2004). Beta distribution depends on two scaling parameters Be(*p, q*) and it can also be parametrized in terms of its mean 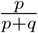, a dispersion parameter *p* + *q*, and the variance 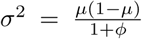. This reparametrization supports the truncated nature of the beta distribution, where the variance depends on the mean and maximum variance is observed at the centre of the distribution whereas it is minimum at the edges. In addition, the dispersion of the distribution, for fixed *µ*, decreases as *ϕ*. The density function is

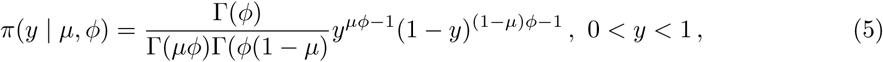

where Γ is the gamma function.

Let *y*_1_, …, *y*_*n*_ be independent beta variables, where each *y*_*i*_, *i* = 1, …, *n*, with mean *µ* and unknown precision *ϕ*. These variables, representing proportions (in our particular case, cumulative proportion of ascospores discharged), can be linked to the linear predictor using a similar approach to the generalized linear models (GLM) with the logit function:

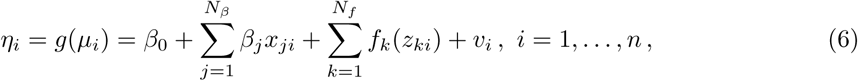

where *η*_*i*_ enters the likelihood through a logit link, *β*_0_ is the intercept of the model, *β*_*j*_ are the fixed effects of the model, *f*_*k*_ denote any smooth effects, and *v*_*i*_ represents unstructured error terms (random variables). The models which we deal with in this work include only fixed effects and in some cases an unstructured term corresponding to independent random effect year, but they could also incorporate spatial or spatio-temporal effects (Paradinas et al., 2018).

In this study we used the approach by Smithson and Verkuilen (2006), who proposed a transformation which compresses the data symmetrically around 0.5. In particular, the transformed values were obtained as

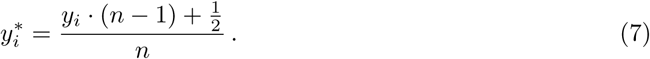

### 2.3 Bayesian inference with INLA

A Bayesian hierarchical approach was used to approximate the variation in the proportion of discharged ascospores with INLA (Rue et al., 2009). This methodology uses Laplace approximations (Tierney and Kadane, 1986) to get the posterior distributions in latent Gaussian models (LGMs) (Rue et al., 2009). LGMs are a particular case of the structured additive regression (STAR) models, where the mean of the response variable is linked to a structured predictor that accounts for the effects of various covariates in an additive way. The prior knowledge of the additive predictor is expressed using Gaussian prior distributions. In this context, all the latent Gaussian variables can be seen as components of a vector known as the latent Gaussian field.

Vague Gaussian distributions *β*_*j*_ *∼𝒩* (0, *τ* = 10^−3^) were used here for the parameters involved in the fixed effects (i.e. *ADD, ADDvpd* and *ADDwet*) and a multivariate independent Gaussian distribution for the random effect year, depending of a precision parameter *v*_*i*_*∼ 𝒩* (**0**, ***Q***(*τ*)). Precision of the beta distribution (*ϕ*) was reparametrized as *ϕ* = exp(*α*) to assure that *ϕ* was a positive parameter. We assumed, following Simpson et al. (2017), pc-priors on the logprecision for both parameters.

The computational implementation R-INLA for R was used to perform approximate Bayesian inference (R Core Team, 2018). Pearson’s correlation coefficients among *ADD, ADDvpd* and *ADDwet* were previously calculated to assist in variable selection and minimize potential problems of multicollinearity. Model selection was conducted based on choosing the best subset of covariates. This method evaluates all 2^*k*^ (where k represents the number of components of the model: covariates and the random effect in our case) possible models and choose the best model according to information criteria (Heinze et al., 2018). In this work we used the deviance information criterion (DIC), which is a generalization of the Akaike information criterion (AIC) developed for Bayesian model comparison (Spiegelhalter et al., 2002), and the Watanable-Akaike information criteria (WAIC) (Watanabe, 2010). The DIC and WAIC are the sum of two components, one quantifying model fit and other evaluating model complexity. The predictive ability of the models was evaluated by cross validation using the logarithmic conditional predictive ordinate (LCPO) (Roos et al., 2011). Models with the lowest values of DIC, WAIC and LCPO were selected.

Lastly, the marginal posterior densities for the parameters and predictive distributions for new observations were obtained with the best model at L’Alcúdia. Median values of the posterior predictive distribution were linearly regressed against the observed values and *R*^2^ was computed. The mean absolute error (MAE), mean square error (MSE) and root mean square error (RMSE) were also calculated. The best model at L’Alcúdia was evaluted at Villanueva de Castellón and Guadassuar (2010 and 2011) and Moncada (2012 to 2015). Likewise, linear regression of predicted *vs*. observed, MAE, MSE and RMSE were calculated in each case.

## 3 Results

During the period of study at L’Alcúdia, annual mean temperature ranged from 16.51ºC in 2010 to 18.24ºC in 2014 (Figure 1a). The lowest daily mean temperature ranged from 1.32ºC in 2009 to 5.38ºC in 2013 and the highest daily mean temperature from 27.79ºC in 2013 to 32.23ºC in 2010. Annual mean relative humidity ranged from 66.49% in 2012 to 72.63% in 2015. The lowest daily mean relative humidity was 28.58 in 2012 and the highest daily mean relative humidity was 98.80% in 2015. The number of days with *vpd ≤* 4*hPa* ranged from 62 in 2013 to 133 in 2015. Annual rainfall ranged from 208.3 mm in 2014 to 698.8 mm in 2009. The number of days with rainfall *≥* 0.2mm ranged from 63 in 2013 to 117 in 2010 and 2011. Annual *ADD* ranged from 6027.95 in 2010 to 6659.18 in 2014 (Figure 1b). Annual *ADDvpd* ranged from 704.73 in 2013 to 1764.47 in 2015 and *ADDwet* from 382.00 in 2013 to 883.51 in 2015.

**Figure 1:**
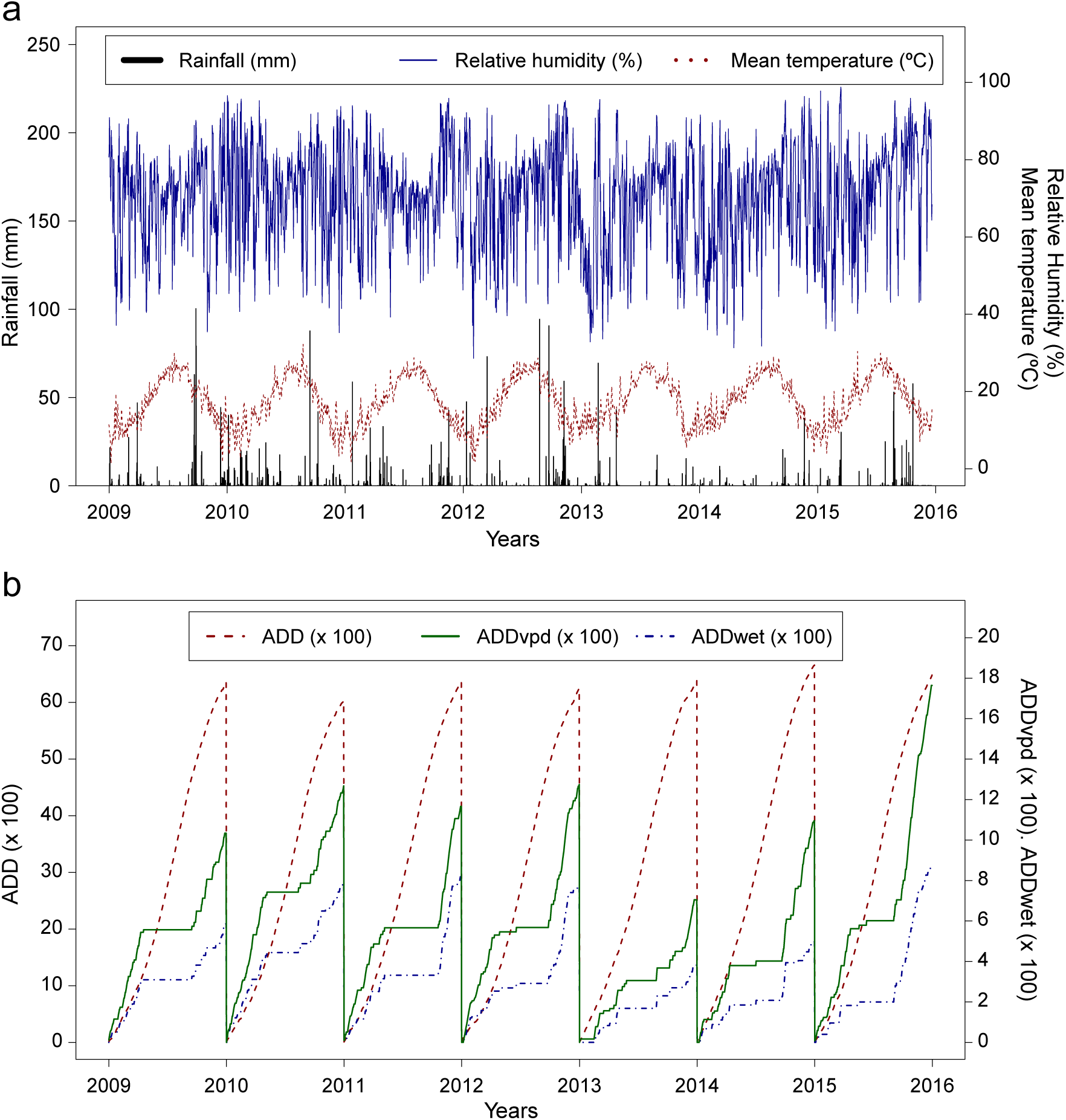
Environmental conditions in the study orchard at L’Alcúdia from 2009 to 2015 a: Rainfall, relative humidity and mean temperature. b: Accumulated degree days (*ADD*), *ADD* considering vapor pressure deficit (*ADDvpd*) and *ADD* considering vapor pressure deficit and rainfall (*ADDwet*).

The best models for the cumulative proportion of *P. nawae* ascospores discharged from persimmon leaf litter are displayed in Table 1. Most of them included the random effect year (***v***) and those not including the fixed effect *ADD* were ranked very low based on their DIC, WAIC and LCPO values. Two of the five best models included all three fixed effects, *ADD, ADDvpd* and *ADDwet*, but were not further considered because the Pearson’s correlation coefficient between *ADDvpd* and *ADDwet* was 0.85, indicating potential problems of multicollinearity. The selected model, having the lowest values of DIC, WAIC and LCPO, included the fixed effects *ADD* and *ADDvpd*, and the random effect year (***v***). Linear regression of the median posterior predictive distribution against observed values accounted for more than 95% of the total variance (*R*^2^ = 0.98), with an intercept of 0.025 and a slope of 0.950, respectively (Figure 3). The MAE for this model was 0.042, the MSE was 0.004 and the RMSE 0.062.

**Table 1:** Models for the cumulative proportion of *Plurivorosphaerella nawae* ascospores discharged from persimmon leaf litter based on accumulated degree-days (*ADD*), *ADD* taking into account vapor pressure deficit (*ADDvpd*), *ADD* taking into account vapor pressure deficit and rain (*ADDwet*), and the random effect year (***v***).

| MODEL<sup>1</sup> | DIC<sup>2</sup> | WAIC<sup>3</sup> | LCPO<sup>4</sup> |
| --- | --- | --- | --- |
| $1 + ADD + ADDvpd + \mathbf{v}$ | -1185.39 | -1183.49 | -1.88 |
| $1 + ADD + ADDwet + ADDvpd + \mathbf{v}$ | -1184.84 | -1183.22 | -1.88 |
| $1 + ADD + ADDwet + \mathbf{v}$ | -1165.41 | -1163.80 | -1.85 |
| $1 + ADD + \mathbf{v}$ | -1155.80 | -1153.73 | -1.83 |
| $1 + ADD + ADDvpd$ | -1117.62 | -1115.66 | -1.77 |
| $1 + ADD + ADDwet + ADDvpd$ | -1115.70 | -1113.80 | -1.77 |
| $1 + ADD + ADDwet$ | -1033.69 | -1032.96 | -1.64 |
| $1 + ADD$ | -970.37 | -970.11 | -1.54 |
| $1 + ADDwet + ADDvpd + \mathbf{v}$ | -619.42 | -621.50 | -0.99 |
| $1 + ADDvpd + \mathbf{v}$ | -609.87 | -611.25 | -0.97 |
| $1 + ADDwet + \mathbf{v}$ | -575.94 | -578.17 | -0.92 |
| $1 + ADDwet + ADDvpd$ | -470.71 | -472.11 | -0.75 |
| $1 + ADDvpd$ | -469.37 | -470.37 | -0.75 |
| $1 + ADDwet$ | -455.96 | -456.94 | -0.72 |
| $1 + \mathbf{v}$ | -393.96 | -395.46 | -0.63 |
| 1 | -375.73 | -376.37 | -0.60 |

<sup>1</sup> $biofix = 1$ January, $T_{base} = 0^{\circ}\text{C}$ .
<sup>2</sup>Deviance information criterion.
<sup>3</sup>Watanabe-Akaike information criteria.
<sup>4</sup>Logarithmic conditional predictive ordinate.

In the selected model, both *ADD* and *ADDvpd* were relevant. The parameter for the fixed effect *ADD* had a mean posterior distribution of 0.293 with a 95% credible interval [0.278, 0.308], not overlapping with zero (Table 2). The parameter for the fixed effect *ADDvpd* had a mean posterior distribution of 0.443 with a 95% credible interval [0.313, 0.575], not overlapping with zero either. The posterior distribution of the two hyperparameters was also computed (Table 2), showing that the random effect had low precision and it was relevant in our model. The two fixed effects, *ADD* and *ADDvpd*, had positive effects on the expected cumulative proportion of *P. nawae* ascospores discharged from the leaf litter, so the cumulative proportion of ascospores increased when *ADD* and *ADDvpd* incremented. Considering the median posterior predictive distribution, 5% of *P. nawae* ascospores were discharged from 995 to 1520 *ADD* and 190 to 545 *ADDvpd* (Figure 2). These thermal times corresponded to 4 April and 25 April. At the other extreme, 95% of *P. nawae* ascospores were discharged from 2585 to 3260 *ADD* and 300 to 740 *ADDvpd* (Figure 2). These thermal times corresponded to 28 June and 30 July. For the 0.025 quantile of the posterior predictive distribution, 5% of ascospores were discharged from 1250 to 1930 *ADD* and 270 to 720 *ADDvpd*, and 95% of ascospores discharged from 3320 to 4180 *ADD* and 300 to 750 *ADDvpd*. For the 0.0975 quantile, 5% of ascospores discharged from 610 to 830 *ADD* and 155 to 250 *ADDvpd*, and 95% of ascospores discharged from 2040 to 2745 *ADD* and 300 to 745 *ADDvpd*.

**Table 2:**
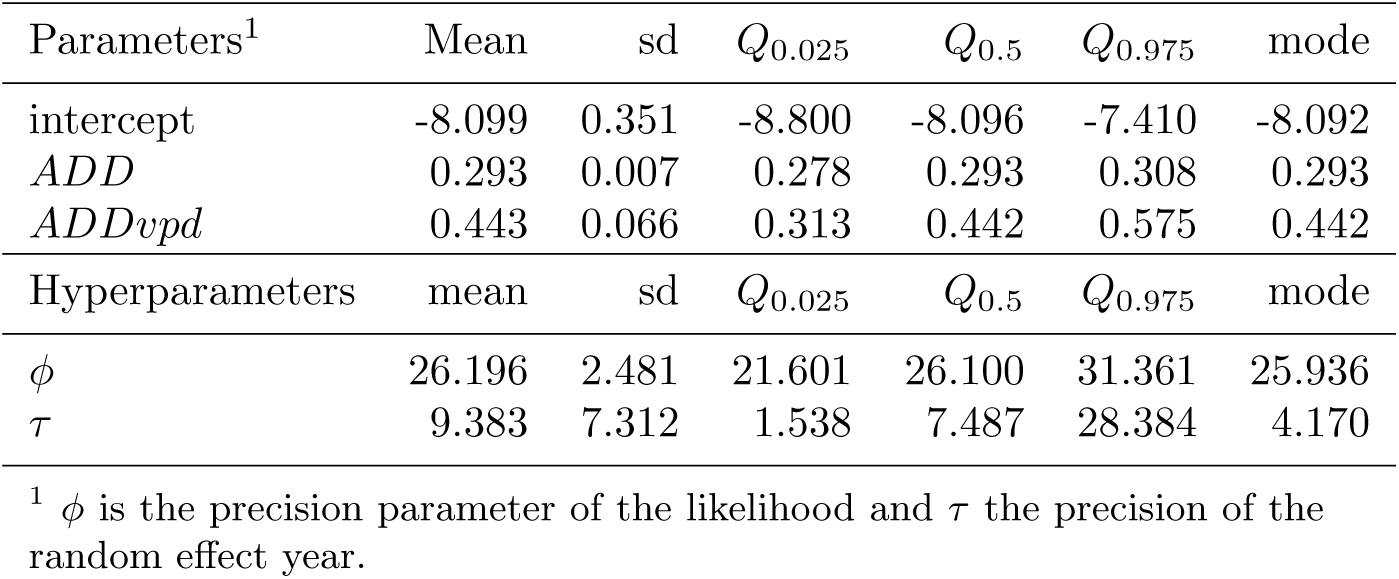
Model for the cumulative proportion of *Plurivorosphaerella nawae* ascospores discharged from persimmon leaf litter including the fixed effects accumulated degree-days (*ADD*) and *ADD* taking into account vapor pressure deficit (*ADDvpd*), and the random effect year. Mean, standard deviation (sd), quantiles (*Q*) and mode for the parameters and hyperparameters (*ϕ, τ*).

**Figure 2:**
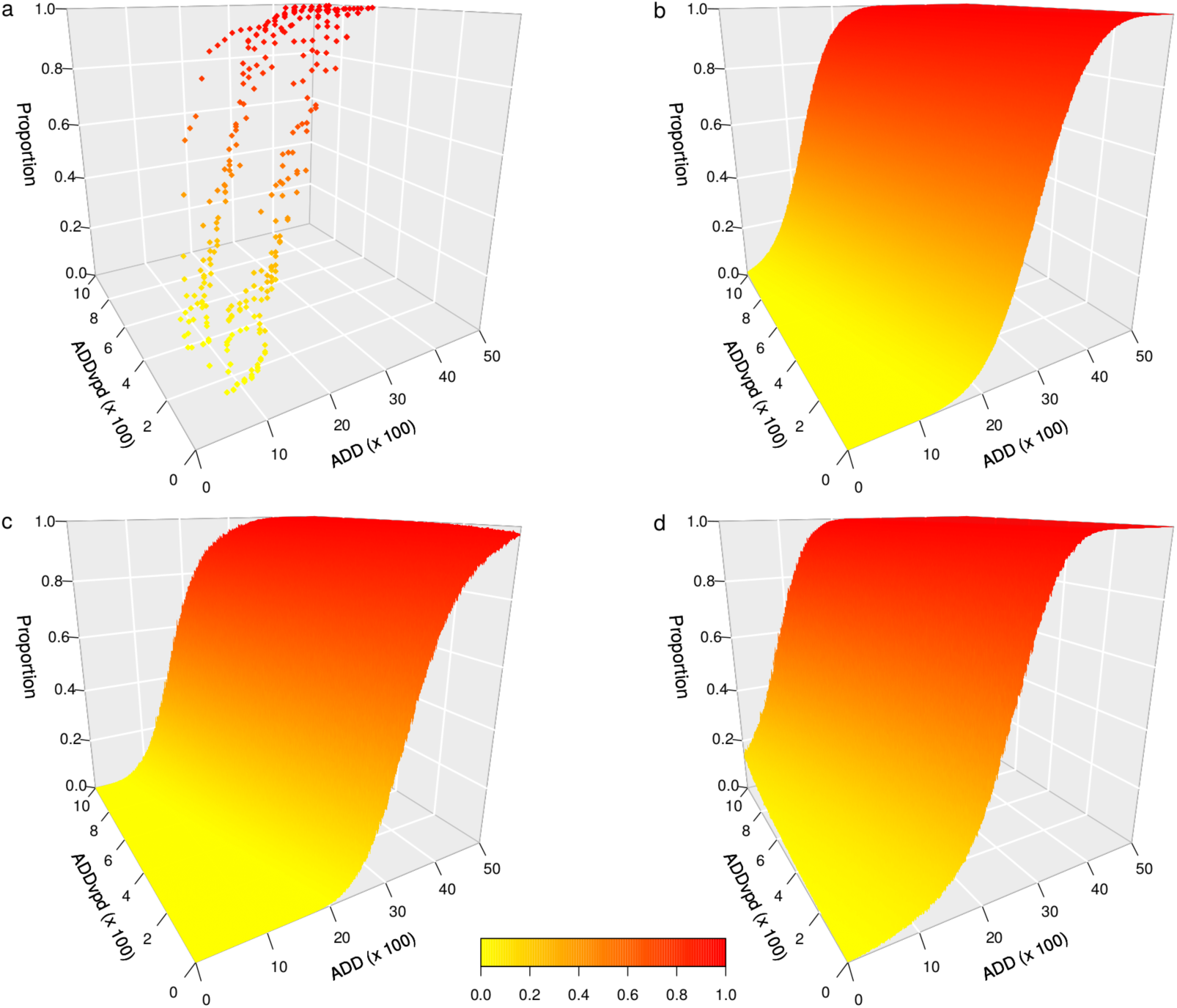
Model for the cumulative proportion of *Plurivorosphaerella nawae* ascospores discharged from persimmon leaf litter at L’Alcúdia based on accumulated degree days (*ADD*) and *ADD* considering vapor pressure deficit (*ADDvpd*). a: data, b: median posterior predictive distribution, c and d 95% credible interval.

**Figure 3:**
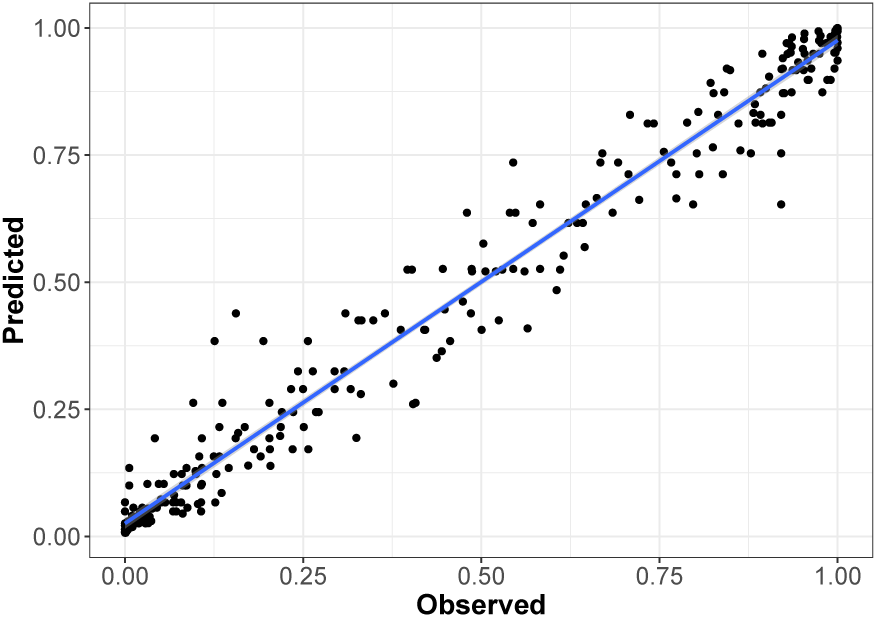
Linear regression between observed values and the median of the posterior predictive distribution for the model of the cumulative proportion of *Plurivorosphaerella nawae* ascospores discharged from persimmon leaf litter (black dots) at L’Alcúdia. Blue line is the regression line.

When the selected model was applied to the evaluation dataset (Table 3), values of MAE > 0.1 were obtained at Moncada in 2012 (0.311), 2014 (0.128) and 2015 (0.275) as well as at Benimodo in 2010 (0.109). A MAE *<* 0.05 was obtained at Guadassuar in 2010 (0.042) and at Moncada in 2013 (0.042). For the RMSE (i.e. 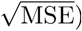, values > 0.3 were obtained only at Moncada in 2012 (0.371) and 2015 (0.334). Values of RMSE *<* 0.1 were obtained at Moncada in 2013 (0.058), at Guadassuar in 2010 (0.062) and at Benimodo in 2011 (0.097). When the median of the posterior predictive distribution was linearly regressed against the observed values, *R*^2^ > 0.90 were obtained but at Moncada in 2012 (0.720) and 2015 (0.700), at Benimodo in 2010 (0.849) and Guadassuar in 2011 (0.892). When plotting observed *vs*. predicted values, these four location/year combinations also showed the poorest graphical fit (Figure 4). In general, residuals were greater during the exponential phase and lower at the beginning and end of the ascospore production period (Figure 4). Considering all the orchards and years of the evaluation dataset, the model predicted 5% and 95% ascospore discharge from 4 April to 24 April and from 27 June to 18 July, respectively.

**Table 3:** Mean absolute error (MAE), mean square error (MSE) and root mean square error (RMSE) for the model of the cumulative proportion of *Plurivorosphaerella nawae* ascospores discharged from persimmon leaf litter at Benimodo, Villanueva de Castellón, Guadassuar and Moncada. Values of *R*^2^ for the linear regression between observed values and the median posterior predictive distribution.

| Location | Year | MAE | MSE | RMSE | $R^2$ |
| --- | --- | --- | --- | --- | --- |
| Benimodo | 2010 | 0.109 | 0.036 | 0.189 | 0.849 |
|  | 2011 | 0.073 | 0.009 | 0.097 | 0.946 |
| V. Castellón | 2010 | 0.072 | 0.014 | 0.119 | 0.934 |
|  | 2011 | 0.070 | 0.013 | 0.114 | 0.972 |
| Guadassuar | 2010 | 0.042 | 0.004 | 0.062 | 0.981 |
|  | 2011 | 0.094 | 0.024 | 0.156 | 0.892 |
| Moncada | 2012 | 0.311 | 0.138 | 0.371 | 0.720 |
|  | 2013 | 0.042 | 0.003 | 0.058 | 0.979 |
|  | 2014 | 0.128 | 0.027 | 0.165 | 0.917 |
|  | 2015 | 0.275 | 0.112 | 0.334 | 0.700 |

**Figure 4:**
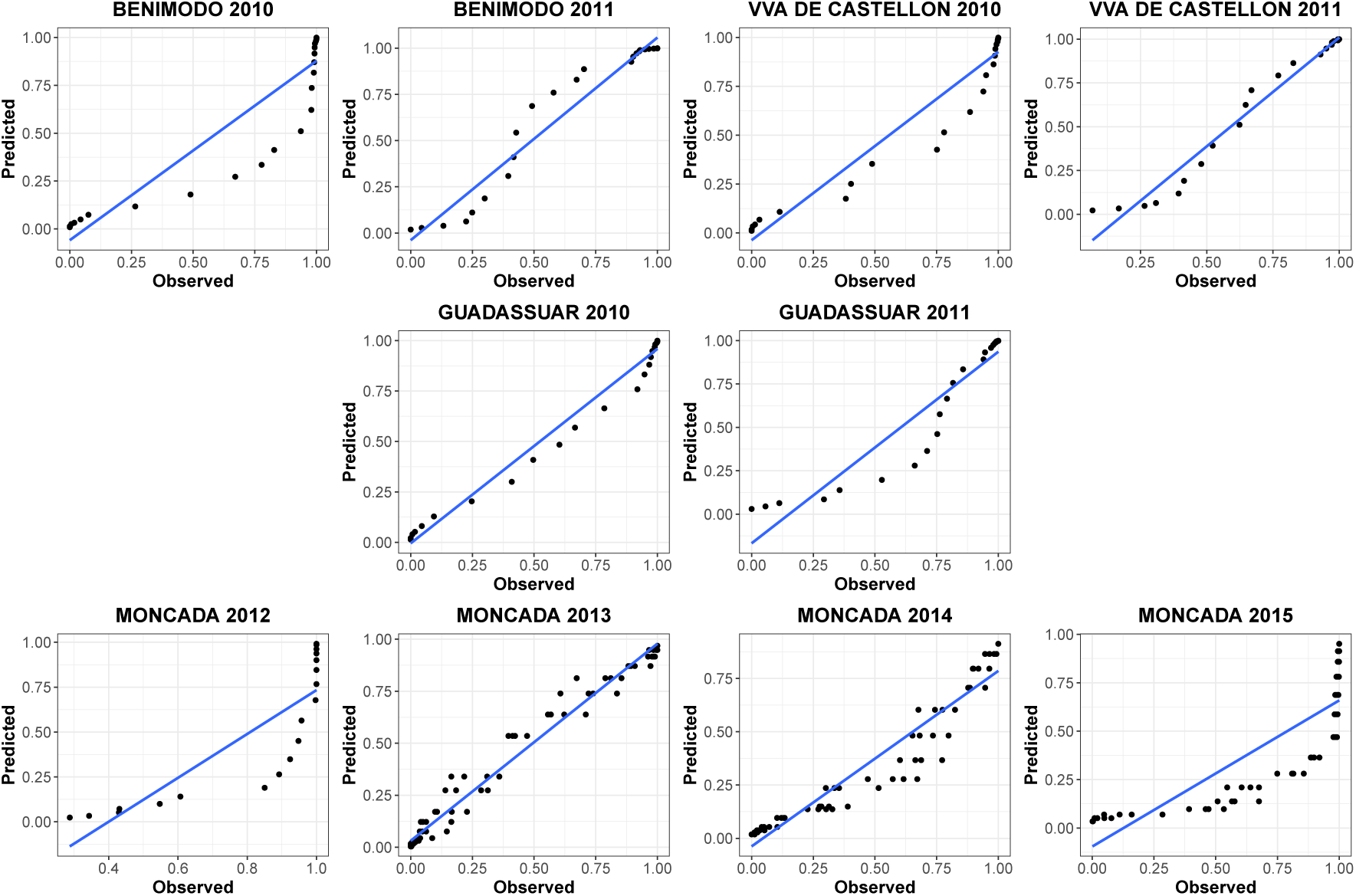
Linear regression between observed values and the median of the posterior predictive distribution for the model of the cumulative proportion of *Plurivorosphaerella nawae* ascospores discharged from persimmon leaf litter (black dots) at Benimodo, Villanueva de Castellón, Guadassuar and Moncada. Blue line is the regression line.

## 4 Discussion

In the present study we propose a Bayesian beta regression framework to model the dynamics of inoculum production when dealing with proportion data. Beta regression overcomes all the drawbacks of the traditional data transformations (Ferrari and Cribari-Neto, 2004). First, it allows a direct interpretation of model parameters in terms of the original data; second, the analysis is not sensitive to the sample size; and lastly, posterior distributions are expected to concentrate well within the bounded range of proportions. Beta regression is widely applied in many scientific disciplines, but in plant pathology it has been used only to a limited extent. Busby et al. (2013) used beta regression to evaluate the effects of fungal endophytes and *Populus* genotypes on the proportion of leaf area affected by *Drepanopeziza populi*. Yellareddygari et al. (2016) developed a beta regression model to predict the incidence of pink rot, caused by *Phytophthora erythroseptica*, on potato tubers during storage based on disease incidence at harvest. Burman et al. (2017) estimated the potential geographic distribution of *Austropuccinia psidii* in Puerto Rico with beta regression. More recently, Xu et al. (2019) related the proportion of wheat plants carrying overwintered *Puccinia striiformis* f. sp. *tritici* with temperature-derived variables using beta regression.

One of the known drawbacks of the beta distribution is its incapability to provide a satisfactory description of the data at the extremes, i.e. 0 and 1 (Ferrari and Cribari-Neto, 2004). Several solutions have been presented in the literature, like adding a small error value to the observations to satisfy this criterion (Warton and Hui, 2011) or using zero and one inflated models (Liu and Kong, 2015). In our study we adopted the approach by Smithson and Verkuilen (2006), who proposed a transformation which compresses the data symmetrically around 0.5, and so, extreme values are affected more than values lying close to 0.5.

Previous studies with beta regression in the context of plant pathology used frequentist inference and did not include random effects (Busby et al., 2013; Yellareddygari et al., 2016; Burman et al., 2017; Xu et al., 2019). In contrast to frequentist inference, where point estimates and confidence intervals are obtained for model parameters, results of Bayesian inference are presented by their posterior distributions. By means of the Bayes theorem, these posterior distributions combine the prior knowledge about the parameters as well as the information gathered from experiments expressed via the likelihood. In frequentist inference, the 100(1 *− α*)% confident interval is defined such that, if the data collection process is repeated again and again, then 100(1 *− α*)% of the confidence intervals formed would contain the unknown parameter value (Fisher, 1956). However, in Bayesian inference, uncertainty of the parameters is typically displayed by their credible intervals. The interpretation of the Bayesian 100 (1 *− α*)% credible interval is that this interval contains 100(1 *− α*)% of the posterior distribution of the parameter, so the probability for the parameter of interest to be in that interval is 1 *− α* (Gelman et al., 2013).

Although the Bayesian approach allows to incorporate prior information of the parameters, as no information was available for *P. nawae*, we used vague priors with large variance reflecting great uncertainty (Carlin and Louis, 2008). The hierarchical structure enabled for a more natural specification of the model, particularly when, as in our case, random effects are included. These complex models can be difficult to solve with frequentist inference. However, they can be readily approached from a Bayesian perspective. Posterior distributions in complex models do not have a closed form and numerical approaches such as MCMC are generally needed to approximate them (Gilks et al., 1996). The INLA approach was used here instead of MCMC because of its higher computation efficiency and speed of calculation, as well as its good behaviour for beta regression models (Rue et al., 2009; Paradinas et al., 2018). Despite its advantages over MCMC, particularly when dealing with large datasets, only a few studies using INLA are available in plant disease epidemiology (Marcais et al., 2016; Martínez-Minaya et al., 2018; Denis et al., 2018).

In the model selected for *P. nawae, ADD* and *ADDvpd* were the covariates driving the maturation of ascospores (Table 2). It was described that *P. nawae* overwinters in the leaf litter as mycelia or pseudothecial primordia, which mature and form ascospores as temperatures raise in spring. Ascospores are then released when pseudothecia absorbe enough moisture (Kwon and Park, 2004; Vicent et al., 2011, 2012). Nevertheless, quantitative relationships between ascospore production and environmental variables were not available for *P. nawae*. There are, however, many examples in the literature for other ascomycetes indicating that models for ascospore maturation should be corrected for dry periods, by accumulating degree-days only when enough moisture was available in leaf litter. Navas-Cortés et al. (1998b) considered only *ADD* on rainy days (*≥* 1mm) to predict the maturation of *Mycosphaerella rabiei* pseudothecia in chickpea in Spain. This study indicated that rain was essential for the synchronization between *M. rabiei* ascospore availability and the vegetative growth of the host. In Norway, Stensvand et al. (2005) improved model accuracy for *V. inaequalis* ascospore maturity in dry years by halting degree-day accumulation if seven consecutive days without rain occurred. When comparing models for *V. inaequalis* ascospore maturation in different areas, Eikemo et al. (2011) indicated that those adjusted for dry periods were consistently the most accurate predictors of ascospore depletion.

Interestingly, *ADDvpd* based on *rh* was more relevant in the model for *P. nawae* than *ADDwet*, which included also the effect of rain (*≥* 0.2 mm) (Table 1). During the period of study, dew resulting from high *rh* was much more frequent than rain (Figure 1a). In the case of *P. nawae*, wetness induced by dew was not sufficient for substantial ascospore discharge (Vicent et al., 2011), but in absence of rain it may favor pseudothecial development and subsequent ascospore maturation. This was described by Rossi et al. (1999) for *V. inaequalis* in Italy, where models accounting for leaf litter wetness significantly improved estimates of airborne ascospores. Furthermore, Mondal and Timmer (2002) demonstrated that alternate wetting and drying of the leaf litter was necessary for the formation of pseudothecia of *Zasmidium citri-griseum*.

The selection of the date from when degree-days are accumulated (i.e. biofix), has been pointed out as a critical factor in the models for ascospore maturation and release. In some cases, a date was chosen based on a specific phenological stage of the host, such as bud break or green tip (MacHardy and Gadoury, 1985; Eikemo et al., 2011). However, the synchrony between host and fungal phenology may differ among regions. Often, the date of detection of the first mature pseudothecia or the first ascospore trapped has been used as the biofix (Spotts et al., 1994; Eikemo et al., 2011). Nevertheless, this approach relies upon the sensitivity of the detection methods used and, more importantly, requires leaf litter sampling or deployment of spore traps. Both methods are time and resource consuming, limiting the extent and density of the monitoring network. The most convenient approach to set the biofix is using a fixed calendar date (James and Sutton, 1982a), but it was argued that it does not take into account the climatic differences between regions (Llorente and Montesinos, 2004). Roubal and Nicot (2016) used numerical optimization to define a single calendar date (1 January) as the biofix for *V. inaequalis*. In our case, 1 January was also chosen as the biofix for *P. nawae* because, in our conditions, persimmon trees attain complete leaf fall around this date and so all the leaves are on the orchard floor still with undifferentiated ascocarps.

Like for other ascomycetes, our model for *P. nawae* considered temperature and moisture covariates as having a continuous positive effect on ascospore development (Figure 2). However, the process resulting in ascospore formation in the leaf litter can be divided in different phases, which may have distinct temperature and moisture requirements. For *M. rabiei*, Gamliel-Atinsky et al. (2005) defined pseudothecium ontogeny followed by initiation of asci and ascospores, and finally ascospore maturation. Navas-Cortés et al. (1998a) indicated that moisture was essential for pseudothecium ontogeny in *M. rabiei* whereas cool temperatures were required for the initiation of asci and ascospores. Actually, low temperatures are generally needed for the onset of sexual reproduction in many ascomycetes (Trapero-Casas et al., 1992; Scherm et al., 2001). James and Sutton (1982b) indicated that the development of asci and ascospores in *V. inaequalis* was initiated in spring, after a dormant period which was not influenced by temperature or moisture levels. Then, rapid maturation of ascospores was favored by moisture and increasing temperatures. Gadoury and MacHardy (1982) indicated that the productivity of *V. inequalis* pseudothecia and the rate of asci maturation were inversely proportional to temperatures from 4 to 20ºC. However, the rate of ascospore maturation was directly proportional to temperature within this range.

Roubal and Nicot (2016) related temperature to ascospore production of *V. inaequalis*, obtaining better results when using a nonlinear unimodal function of thermal time compared with *ADD*. This unimodal function accounted for reduced effects of low and high temperatures on ascospore production. Actually, this type of unimodal response to temperature was reported for some ascomycetes and ectotherms in general (Naseri et al., 2008; Trudgill et al., 2005). Nevertheless, the relationship between the rate of development and temperature is often linear over much of the range up to the thermal optimum, and thus *ADD* are usually considered for thermal time calculations (Trudgill et al., 2005). In any case, knowledge about the temperature and moisture requirements for each phase of ascospore formation in *P. nawae* may help to develop models with improved performance and better extrapolation to other areas.

Our models also corroborated previous studies in Spain indicating that *P. nawae* adapted to semi-arid conditions by advancing the period of ascospore production to escape from the typical Mediterranean rain-less summer. Consequently, ascospore production coincides with rains in spring, from March to June, under more favorable conditions for infection. On the other hand, low winter temperatures in Korea delayed ascospore release to June-August, then synchronized with the abundant summer rains typical in this area (Kang et al., 1993; Kwon et al., 1995; Kwon and Park, 2004).

In previous studies, discharge tests allowed detection of mature ascospores of *P. nawae* in the leaf litter before they were released to air in the orchard (Vicent et al., 2012). Similar results were reported for *Sphaerulina musiva* in poplar, where peak ascospore production in leaf litter measured with discharge tests occurred 7 days earlier than peak airborne ascospores (Luley and McNabb Jr, 1991). However, when comparing different methods to estimate the maturity and release of *V. inaequalis* ascospores, Gadoury et al. (2004) found that cumulative ascospore release in discharge tests from the leaf litter lagged behind that measured in the orchard air by spore traps. This was mainly attributed to litter decay, which progressively reduced the leaf litter area on the orchard floor and subsequently the overall ascospore population in the air (Gadoury and MacHardy, 1982; Gadoury et al., 2004). This time lag may be even larger when a fixed leaf area sample instead of a number of leaves is used in discharge tests. In contrast to apple leaves, persimmon leaves are typically coriaceous and no substantial degradation of the leaf litter was observed under the conditions of our study. Indeed, discharge tests from the leaf litter are effectively used by advisory services in Spain to predict airborne inoculum availability and schedule fungicide sprays for *P. nawae* management.

Models for ascospore maturation are mainly aimed to predict the duration of the period for primary inoculum, when fungicide applications need to be intensified. Thus, practical performance of these models relies on their ability to accurately predict ascospore onset and depletion more that the exponential phase of ascospore production (Gadoury et al., 2004; Eikemo et al., 2011). In the case of *P. nawae* in Spain, no secondary conidia have been observed and infections were caused by ascospores formed in the leaf litter (Vicent et al., 2012). Therefore, accurate predictions of the beginning and end of the ascospore production period are paramount for designing efficient fungicide spray programs. Interestingly, when the beta regression model for *P. nawae* was evaluated in different orchards, higher accuracy was obtained at the onset and depletion of ascospore production compared with the exponential phase (Figure 4). Based on our results, we proposed the operating thresholds 5% and 95% of ascospores discharged in a decision support system to schedule fungicide sprays. In the *P. nawae* model, these corresponded with 995-1520 *ADD* and 190-545 *ADDvpd* for the 5%, and 2585-3260 *ADD* and 300-740 *ADDvpd* for the 95% (Figure 2). A test version of the decision support system for CLS control was implemented in the online platform ‘gipcaqui’ from IVIA at http://gipcaqui.ivia.es/avisos-mycosphaerella.

Here a hierarchical Bayesian beta regression was used to model the cumulative proportion of *P. nawae* ascospores produced in persimmon leaf litter. Operating thresholds were proposed for a decision support system to assist advisory services and persimmon growers in optimizing fungicide sprays programs for CLS control in Spain. This same modelling framework can be extended to other ascocmycetes and fungal plant pathogens in general as long as inoculum dynamics are expressed as proportion data.

## Acknowledgements

DC and ALQ would like to thank the Ministerio de Educación y Ciencia (Spain) for financial support (jointly financed by the European Regional Development Fund) via Research Grant MTM2016-77501-P and TEC2016-81900-REDT. JM-M would like to thank for the support by the Basque Government through the BERC 2018-2021 program and by the Ministry of Science, Innovation and Universities: BCAM Severo Ochoa accreditation SEV-2017-0718. AV would like to thank the Instituto Nacional de Investigación y Tecnologíia Agraria y Alimentaria (INIA), Ministerio de Ciencia, Innovación y Universidades via Research Grant RTA2013-00004-C03-02.

